# Development of a KDM6b shRNA conditional knock down mouse model

**DOI:** 10.1101/605618

**Authors:** Julia E. Prier, Marnie E. Blewitt, Ross A. Dickins, Stephen J. Turner

**Affiliations:** The Department of Microbiology and Immunology, at The Peter Doherty Institute, The University of Melbourne, Parkville, Victoria, 3000, Australia; The Epigenetics and Development Division, The Walter and Eliza Hall Institute of Medical Research, Parkville, Victoria, 3052, Australia and The Department of Medical Biology, The University of Melbourne, Parkville Victoria, 3000 Australia; Australian Centre for Blood Diseases, Monash University, Melbourne, Victoria, 3004, Australia; The Department of Microbiology, Biomedicine Discovery Institute, Monash University, Clayton, Victoria, Australia

## Abstract

Antigen-specific CD8^+^ T cell differentiation in response to infection is associated with specific changes in the chromatin landscape resulting in acquisition of the lineage-specific effector functions required for pathogen clearance. Lysine (K)-specific demethylase 6B (KDM6b) is a histone demethylase that specifically recognizes and removes methyl groups from K27 tri/dimethylation on histone 3. This histone modification is associated with a repressive transcriptional state, or, in combination with the active H3K4me3 mark, a bivalent epigenetic state. Resolution of bivalency at fundamental transcription factor loci has been shown to be a key mechanism for the initiation of CD8^+^ T cell differentiation. To begin to address the role of KDM6b in regulating H3K27me3 demethylation in CD8^+^ T cell responses to infection, a model whereby KDM6b levels can be modulated is needed. To address this, we developed a conditional short hairpin RNA (shRNA) mouse model targeting KDM6b. Here we demonstrate that KDM6b knockdown results in diminished naive, CD4^+^ and virus-specific CD4^+^ and CD8^+^ T cell response in response to influenza A infection. To address the molecular mechanism, we demonstrate that KDM6b knockdown resulted in reduced H3K27me3 removal from the *Tbx21* bivalent promoter, compared to luciferase hairpin controls. Surprisingly, this did not necessarily impact T-BET expression, or resolution of other bivalent transcription factor promoters. These data suggest that KDM6b knockdown resulting in diminished IAV-specific CD8+ T cell responses may reflect a demethylase independent function.

## Introduction

Cellular differentiation and fate commitment is mediated by the spatial and temporal control of cell-specific gene expression. While transcription factors mediate the lineage specific expression of genes, alterations to the chromatin structure must first occur to allow the transcriptional machinery to access the DNA. Initiation of gene transcription requires extensive remodelling of chromatin into a structure that promotes transcriptional activation. For example, this may include increased accessibility, addition of H3K4me3 and removal of H3K27me3.

When the cell receives external differentiation signals, the subsequent chromatin remodelling involves enzymes that mediate the addition or removal of histone post translation modifications to establish a transcriptionally permissive chromatin landscape at differentiation-associated genes. Lysine (K)-specific demethylase 6B (KDM6b) is a histone demethylase that specifically recognizes and removes methyl groups from H3K27 tri/dimethylation (1–4). Methylation of K27 at histone H3 is functionally associated with transcriptional repression and gene silencing and hence, its removal is required to permit transcription. The JMJD catalytic domain of Kdm6b catalyses the removal of methyl marks from H3K27 via a Fe2+-dependent dioxygenase mechanism (5, 6).

Several key studies have demonstrated a role for KDM6b in cellular differentiation and fate commitment in embryonic stem cells (ESCs) via removal of H3K27me3 at key lineage specifying genes (7–9). This role for KDM6b has been extended to more mature cell populations such as macrophages and lymphocytes (2). The targeting of KDM6b to specific gene loci appears to be linked via interactions with [lineage-determining?] transcription factors such as RUNX2, and KDM6b plays a role in macrophage activation by interaction with the transcription factor IRF4 (10).

In CD4^+^ T lymphocytes, the differentiation of naïve T_H_0 cells to T_H_1 cells correlates with a decrease in H3K27me3 deposition and consequently the resolution of bivalency at the *Tbx21* locus(11). Interestingly, a recent study showed that knock down of Kdm6b *in vivo* prevented Th1 differentiation (12). However, this did not appear to be due to a lack of resolution of the *Tbx21* locus or a dramatic reduction in *Tbx21*-encoded T-bet expression but rather through altered KDM6b interactions with upstream co-factors (12). Further, the Weinmann group demonstrated a physical interaction of T-bet with KDM6b at the *Ifng* locus *in vitro* in primary CD4^+^ T cells (13). In all these data suggest an essential role for KDM6b in CD4^+^ T cell lineage fate commitment and functional differentiation.

When a naïve CD8^+^ T cell differentiates into an effector CD8^+^ T cell, dramatic epigenetic changes occur to allow for the expression of lineage specific effector function (14–17). As seen upon naïve CD4^+^ T cell activation, antigen-specific CD8^+^ T cell differentiation is associated with the rapid resolution of bivalency at fundamental transcription factor loci to a permissive epigenetic signature (16). What remains to be determined is whether KDM6b dependent H3K27 demethylation is a key step for initiating CD8^+^ T cell differentiation.

Ideally, to be able to fully dissect the role that KDM6b plays during distinct stages of virus-specific CD8^+^ T cell differentiation, a mechanism by which KDM6b levels can be regulated at distinct stages is required. The tetracycline (Tet) inducible system involves use of two distinct genetic elements to regulate target gene expression (18, 19). It requires the generation of two lines of transgenic mice where one contains a tet response element (TRE) promoter that controls specific mRNA or short hairpin RNA (shRNA) expression, and another expressing a tet-transactivator that can activate the TRE promoter. Crossing transgenic mice with tissue-specific expression of the tTA (tet-off) transactivator with mice harboring TRE-driven shRNA allows regulatable knockdown of specific endogenous genes. (20–22). This study describes the generation of transgenic mice that express an inducible KDMB6b shRNA. These mice were then crossed with a variety of tTA-transgenic lines enabling either constitutive or inducible shRNA knockdown. We utilised these mouse lines to initially probe the role of the KDM6b enzyme in regulating CD8+ T cell differentiation in response to infection, with these mice providing a novel resource to study KDM6b knockdown in a variety of other cellular contexts.

## Materials and Methods

### Mice, viruses and infection

Mice (6-10 weeks old) were bred and housed under specific pathogen free conditions in either; the Department of Microbiology and Immunology (DMI) Animal Facility, The Peter Doherty Institute Bioresources Facility (University of Melbourne, Parkville, VIC, Australia) or The Walter and Eliza Hall Institute (WEHI; Parkville, VIC, Australia). All experiments were conducted according to guidelines specified by the university animal ethics committee. For primary influenza A virus (IAV) infection, mice were anaesthetised by isoflourane inhalation at 2% in oxygen and infected intranasally (i.n.) with 10^4^ plaque-forming units (p.f.u.) of A/Hong Kong X31 (A/HKx31, H3N2) in 30 µl of PBS. For secondary infections, mice were first primed intraperitoneally (i.p.) with 1.5×10^7^ pfu mouse adapted A/Puerto Rico/34/8 (A/PR8, H1N1) in 500 µL of PBS at least 6-weeks prior to i.n. challenge with 1×10^4^ pfu HKx31 virus.

### Generation of shRNA retroviruses

Retroviral stocks encoding shRNAs designed to target *Kdm6b* (Table 1) were generated via transfection of 293T cells with the LMP-EBFP2 (21, 23) plasmid harbouring Kdm6b shRNAs, plus plasmids encoding VSVG and PAM-E. The LMP-EBFP2 viral construct enables selection of transduced cells with puromycin and visualisation via EBFP2. Viral supernatants were harvested up to 54 hrs after transfection were used to transduce GP+E86 cells to establish retroviral producer cell lines or pooled and filtered for direct transduction of EL4 cells. Transduction of cells was performed in the presence of 1 mg/ml polybrene (Sigma). Transduced cells were selected for in the presence of 2 µg/ml puromycin (R&D Systems Inc, Minneapolis, MN, USA), live cells harvested by ficoll gradient and flow cytometry used to determine purity based on expression of eBFP indicative of transduction.

**Table 1.**
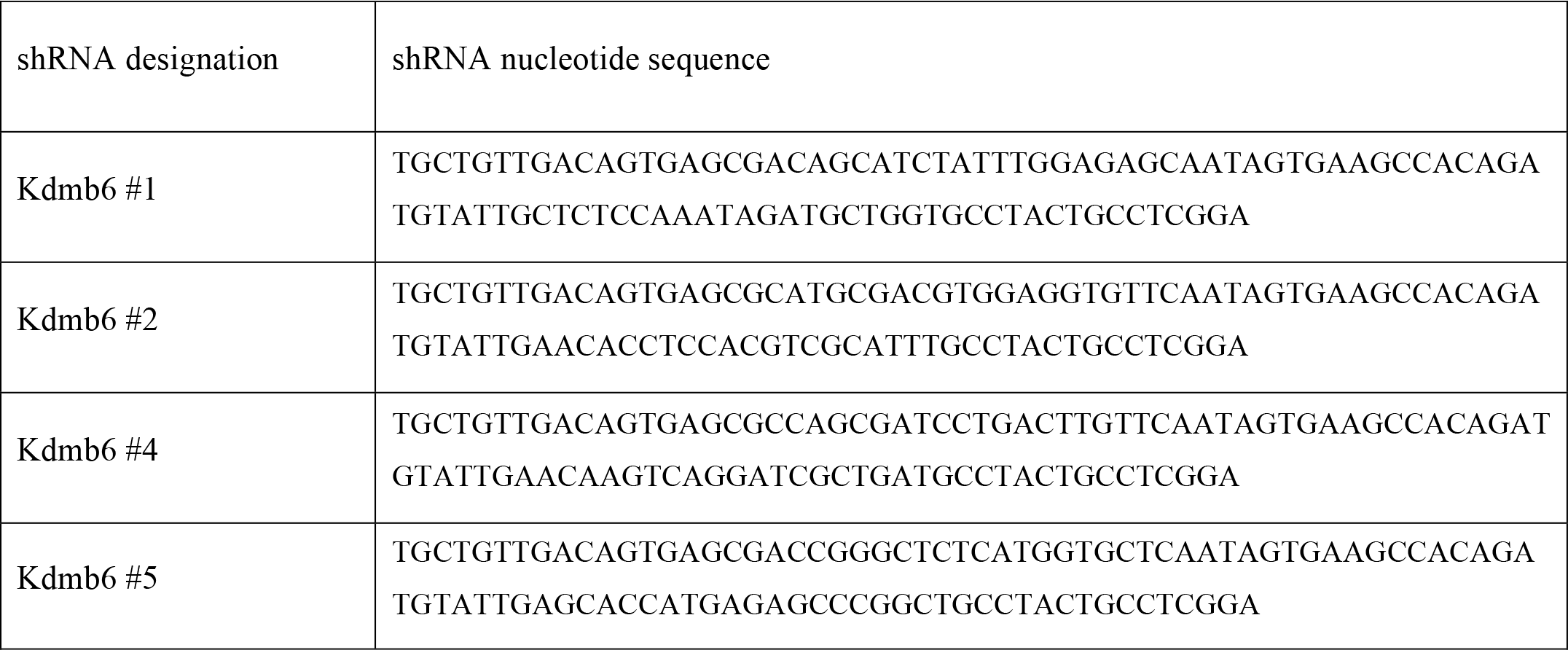
shRNA nucleotide sequences used for in vitro Kdm6b knock-down screen.

To determine knock-down efficiency of selected *Kdm6b* shRNA candidates, 293T or EL.4 cell lines were transduced with titrated levels of pooled supernatants from transfected 293T cells, or supernatant collected from producer cell lines. A population of cells expressing 10-20% eBFP (approximating a low multiplicity of infection) were selected, live cells harvested by ficoll gradient and level of *Kdm6b* knockdown determined by real-time RT-PCR and western blot and compared to non-silencing control.

### Generation of shRNA transgenic mice

Effective shRNA sequences identified using the LMP-EBFP2 vector were excised with XhoI/EcoRI and cloned into a variant of the pCol-TGM vector (24), linking the shRNA to expression of green fluorescent protein (GFP) under control of the TRE3G promoter. This construct was targeted to the *Col1a1* locus in mouse ES cells as described in (24), and these cells were used to generate KDM6b^shRNA^ transgenic mice. Mice were then crossed onto the CMV-rtTA or Vav-tTA transgenic mouse strains (25) to generate KDM6b^shRNA^ × CMV-rtTA or KDM6b^shRNA^ × Vav-tTA mouse lines. As controls, transgenic mice encoding the shRNA for luciferase (Luc^shRNA^) (24) were used for crosses onto CMV-rtTA or Vav-tTA transgenic lines.

### RNA extraction, reverse transcription and RT-PCR

Purified cell were lysed in 0.5–1 ml TRIzol reagent (Invitrogen) for 5 min at RT. RNA was extracted using Direct-zol™ RNA MiniPrep (Zymo Research, Irvine, CA, USA) and if samples were used for RNA-sequencing (RNAseq), were DNase treated with 20 U DNase I (Roche) in a final volume of 50 µl with 1× DNase buffer and 10 U of RNase OUT (Invitrogen) for 20 mins at RT. RNA was then purified and concentrated using QIAgen RNeasy MinElute kit (Qiagen, Hilden, Germany). For reverse transcription, RNA was quantified using a NanoDrop 3300 spectrophotometer (ThermoFisher Scientific) and cDNA synthesized using the Omniscript RT kit (Qiagen) with 2 µM Oligo(dT)_15_ primers (Promega) and 10 U RNase OUT in a 20 µl reaction at 37ºC for 90 min. Real-time PCR was conducted using 1× FAM labelled Taqman^®^ Gene MGB probes with 1-5 ng of cDNA template using 1× Taqman Universal PCR Master Mix (Applied Biosystems). Reactions and analysis were performed in a CFX-Connect Real-Time System with the following cycle conditions; 95ºC 2 min, 40 cycles of 95ºC 15 secs and 60ºC for 1 min. Each reaction was performed in duplicate and results reported as the mean of replicates, normalized to reference gene *L32*, and calculated relative to the indicated samples using the 2^−ΔΔCt^ method (26).

### Flow cytometry and analysis of the IAV-specific CD8^+^ T cell response

Cell suspensions transduced with shRNA lentiviruses were harvested and run on a BD FACSCanto II or BD Fortessa flow cytometer (all BD Biosciences) and data analysed with FlowJo software (Tree Star, Ashland, OR, USA). Mean fluorescence intensity (MFI) was determined as the mean within the positive population. Un-stimulated, isotype control or fluorescence minus one (FMO) control samples were subtracted from test samples as indicated. For analysis of IAV-specific CD8+ T cell populations, splenocytes and cells from bronchoalveolar lavage were stained either CD45.1-PE and CD8-APC, or with CD8-APC and D^b^NP_366_-PE (27) and D^b^PA_224_-PE tetramers(28), at 4°C for 30 minutes and data acquired by FACSCantoII. Gating strategy involved subsequent gating as follows; live cells, singlets, non-auto fluorescent (when possible), non-dump (when possible), lymphocytes followed by gating as indicated in results.

### Chromatin immunoprecipitation

Purified cell suspensions were fixed at indicated concentrations with formaldehyde (Sigma) for 10 min at RT then quenched with 125 mM glycine (Chem Supply, Gillman, SA, Australia) for 10 min at RT. Cells were washed twice with Dulbecco’s PBS (D-PBS) (Gibco) and resuspended at ~4×10^6^ cells/250 µl ChIP lysis buffer prior to sonication using a 130-Watt Ultrapanasonic Processor (Cole-Parmer, Vernon Hills, IL, USA) at 70% amplitude. DNA was sheared into 200-500 bp fragments via a sonication program of 5 secs on, 15 sec rest for a total of 8.5 mins. Pelleting to remove debris was followed by resuspension to the equivalent of 2.5–10×10^5^ cells/ml in ChIP dilution buffer. Samples were pre-cleared with 35 µl Protein A Agarose/Salmon Sperm DNA beads (Millipore, Temecula, CA, USA) at 4ºC for 30 min with continuous rotation. chromatin was immunoprecipitated with antibodies for H3K4me3, H3K27me3 and total H3. Protein-chromatin complexes were purifed with Protein A Agarose/Salmon Sperm DNA beads for 1 hr at 4ºC followed by two sequential washes each of; low salt buffer, high salt buffer, LiCl buffer and TE buffer. Samples were then eluted for 30 min at RT with rotation using 400 µl elution buffer. Eluate were treated with 0.2 M NaCl (AnalaR, Kilsyth, VIC, Australia) overnight at 66ºC to reverse crosslinking and RNA digested with 20 µg RNaseA (Roche) at 37ºC for 1 hr followed by treatment with 20 µg proteinase K (Promega, Madison, WI, USA) at 45ºC for 1 hr to remove protein. DNA was then extracted with an equal volume of phenol:chloroform:isoamyl (25:24:1) (Sigma) and precipitated on ice in 2.5× vol. 100% ethanol (Chem-Supply), 1/10 vol. 3 M NaAc (Sigma) and 1 µl GeneElute (Sigma) overnight. DNA was washed once with 80% ethanol and resuspended in 50-100 µl of 0.1× TE buffer prior to subsequent analysis. Samples were probed by quantitative PCR using primers designed to target specific gene loci (Table 2).

**Table 2.**
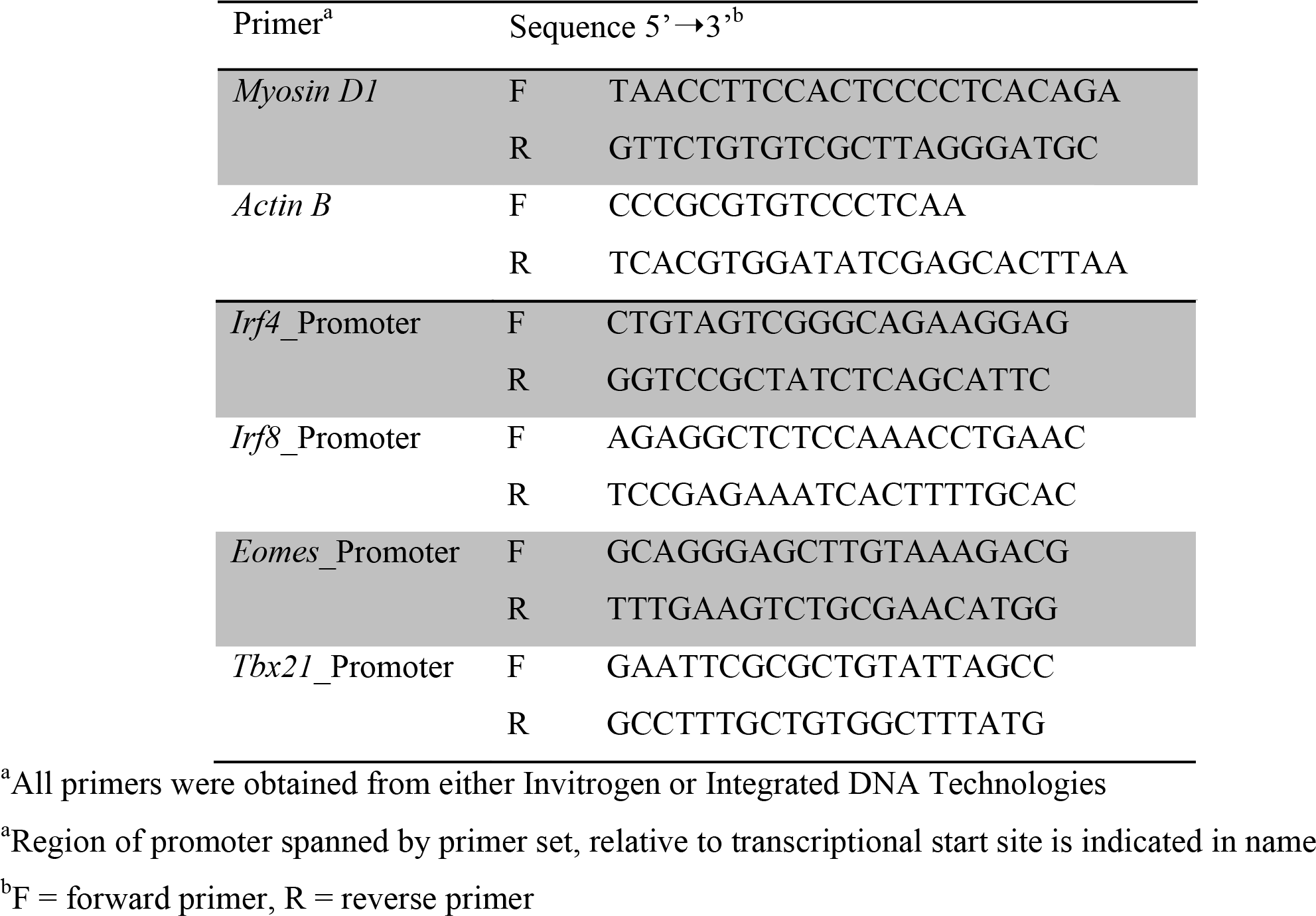
Primers used for SYBR green real-time PCR analysis of ChIP assays.

## Results

### Development of an inducible and reversible Kdm6b knockdown mouse

The tetracycline (tet)-responsive system can be used to reversibly knock down endogenous gene expression by RNA interference (RNAi) (20, 21, 24, 25). To generate shRNA transgenic mice, we first tested a panel of hairpins against KDM6b (Table 1). shRNA expression was linked to fluorescent reporters, enabling easy tracking of cells. 3T3 cell lines were transduced with a series of lentiviral vectors containing different shRNAs each designed to target KDM6b. Efficiency of transduction at low MOI and selection post transduction was determined by enhanced blue fluorescent protein (eBFP) expression (**Fig. 1a**). As early as day one post selection the majority of shRNA transduced 3T3 cells were eBFP^+^ and this remained relatively stable until harvest on day four (**Fig. 1a**). Knockdown efficiency was determined by qRT-PCR analysis of target gene transcription. *Kdm6b* shRNA-1 and 2 resulted in ≥ 70% knockdown compared to non-silencing control shRNA (**Fig. 1b**).

**Figure. 1.**
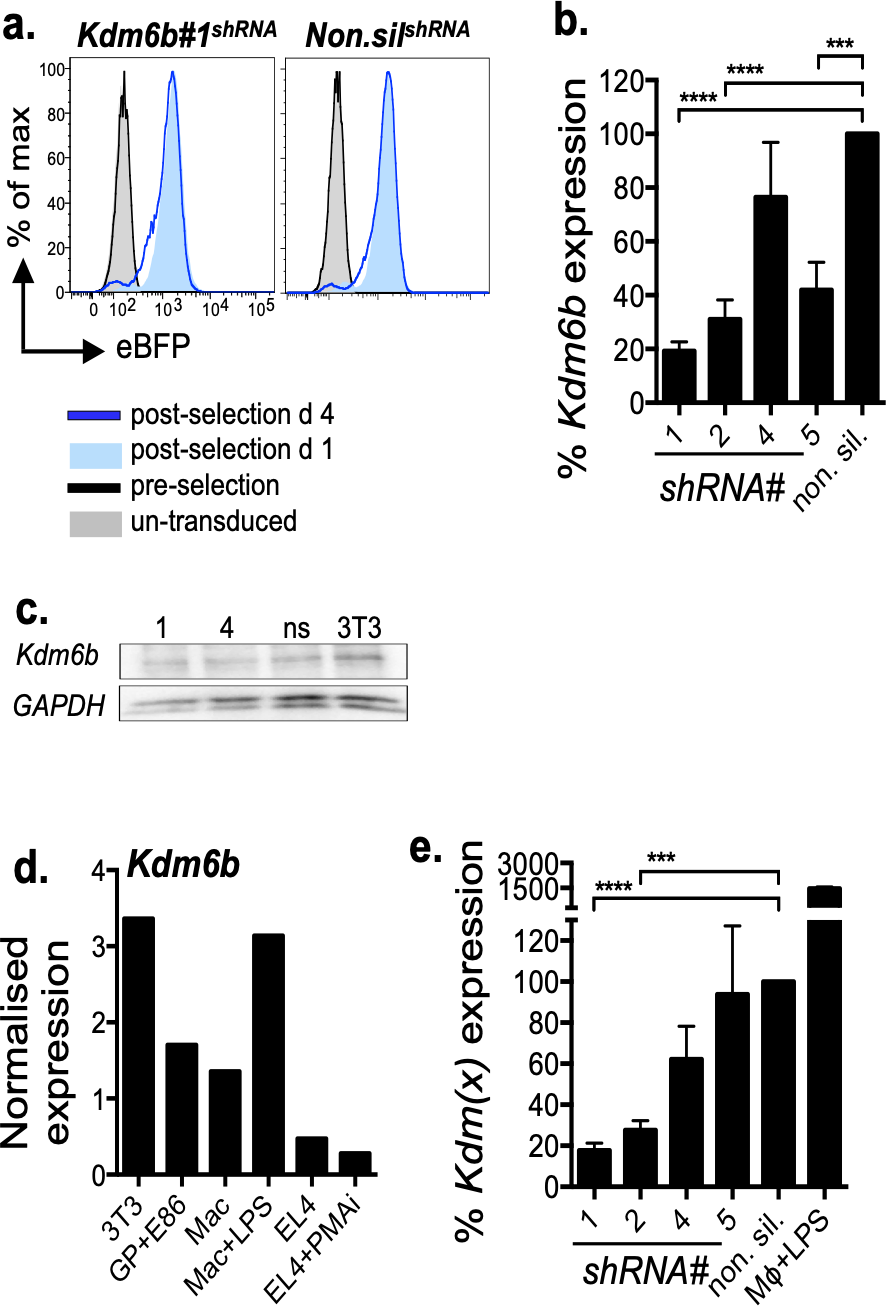
Validating shRNA panels against KDM6b in vitro. 3T3 cells were transduced with lentiviral-shRNA at MOI~1 and selected using puromycin. Flow cytometry plots demonstrate efficiency of selection as determined by eBFP expression (a). Four days following selection, cells were ficoll purified, RNA was extracted with TRIzol reagent, cDNA generated and subjected to qPCR for *L32*, and *Kdm6b* with Taqman primer probes. PCR results were normalised to L32 and the relative gene expression of *Kdm6b* was determined and shown relative to the non-silencing control (b). To determine protein knockdown, lysates were collected following purification as above and probed with indicated antibodies (c). Murine cell lines were probed for *L32* and *Kdm6b* as described in b. at the resting state or following 4 h stimulation with indicated mitogen (d). EL4 cells were transduced with lentiviral-shRNA as described in a. and *Kdm6b* mRNA knockdown was determined as described in b. (e). Data shown are mean+sem and are representative of 3-6 independent experiments or preliminary (d), *p<0.05, **p<0.01, ***p<0.001, ****p<0.0001.

To determine protein expression of the target genes, western blots were performed. *Kdm6b* shRNA-4 resulted in the lowest protein expression compared to non-silencing controls (**Fig. 1c**).

To further validate *Kdm6b* knock down in other cellular contexts, multiple cell lines were tested for their expression levels of *Kdm6b* (**Fig. 1d**). Despite the relatively low expression of *Kdm6b* in EL4 cells, validation was performed in this cell line because it is a C57BL/6 derived mouse T cell lymphoma cell line compared to a fibroblast cell line. Knockdown efficiency of *Kdm6b* was similar in EL4 cells as seen in 3T3 cells (**Fig. 1e**). Kdm6b shRNA-1 had the most efficient and consistent knockdown over both cell types tested and was therefore chosen for continued development into an *in vivo* model.

### Generation of Tetracyline transactivator transgenic mice for Kdm6b shRNA transcription

The expression of shRNA requires robust expression of a tetracycline transactivator (TA) protein. We have previously shown that even under the control of vigorous, cell ubiquitous promoters, there is heterogeneity in both the cell type specificity and level of tet-regulated shRNA transcription. expression (25). Importantly, we demonstrated that effector, effector memory and central memory CD4 and CD8+ T cell subsets showed the highest levels of shRNA expression when the transactivator was under the control of the CAG promoter (CAG-rtTA3; (25)). Therefore, the CAG-rtTA3 transgenic line was selected as the optimal transactivator system to assess the role of KDM6b in activated and virus-specific T cell subsets.

Transgenic *Kdm6b* shRNA or control shLuc1309 (targeting *Luciferase, Luc*) transgenic mice were crossed with CAG-rtTA3 transactivator mice to generate F1 offspring that contain a copy of both the shRNA and the tet-on transactivator (**Fig. 2a**). To validate the knockdown *in vivo*, murine embryonic fibroblasts (MEFs) were generated from F1 offspring and treated with doxycycline for 24 h for analysis of GFP by flow cytometry. Two (of 7) MEF lines isolated from the *Luc*^*shRNA*^.CAG-rtTA cross were GFP^+^ (**Fig 2b**). To confirm that *Kdm6b* knockdown was a reflection of RNA levels, qRT-PCR was performed on GFP^+^ MEFs described above. Importantly, *Kdm6b* mRNA levels were equivalent in untreated *Luc*^*hRNA*^ and *Kdm6b*^*shRNA*^ MEFs (**Fig. 2c**). Upon doxycycline treatment, *Kdm6b* expression was significantly reduced in *Kdm6b*^*shRNA*^ MEFs compared to *Luc*^*shRNA*^ control MEFs (**Fig. 2c**).

**Figure 2.**
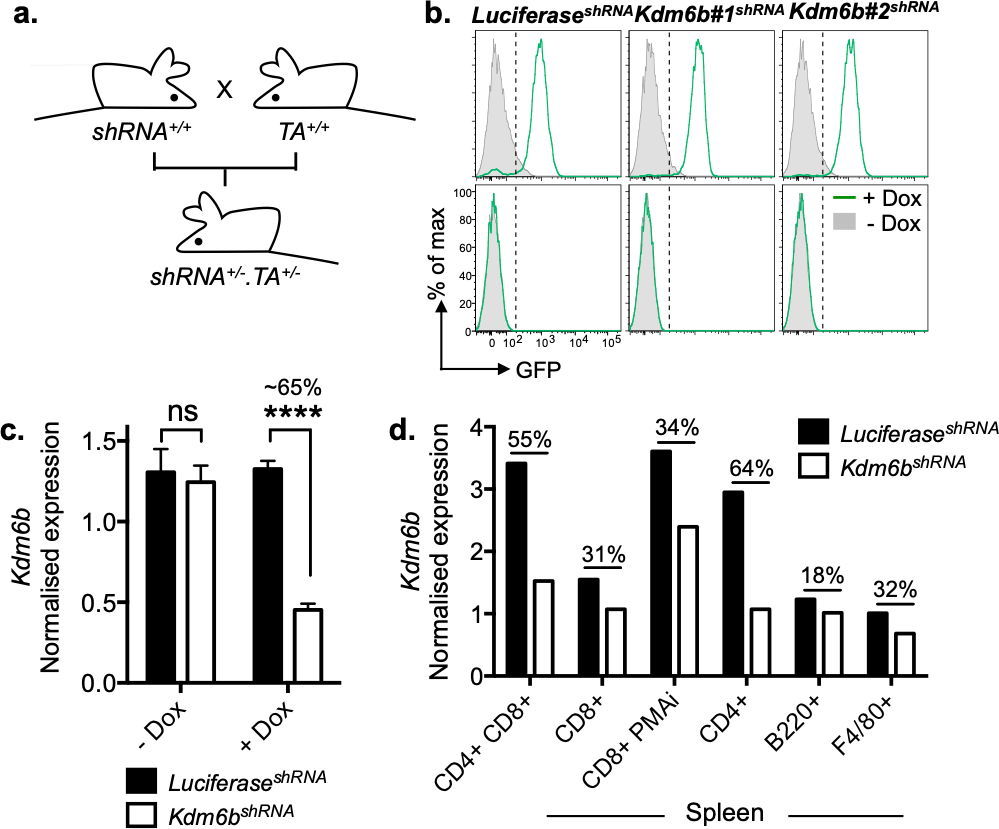
Validating shRNA knockdown of *Kdm6b in vivo*. Breeding strategy to generate F1 offspring (a). GFP reporter expression in MEFs from *Luc*^*shRNA*^. CAG-rtTA and *Kdm6b*^*shRNA*^.CAGrtTA. Top row are representative MEFs containing a copy of both the CAG-rtTA and *shRNA*, bottom row are representative MEFs lacking ability to express hairpin (b). Green histograms indicate cells that received doxycycline and grey histograms are control cells that did not receive doxycycline. MEFs were probed for *L32* and *Kdm6b* as described. Shown are MEFs from *Luc*^*shRNA*^ or *Kdm6b*^*shRNA*^ dams that were GFP+ (white bars, +Dox) and GFP-(black bars, -Dox) (c). Indicated cell types were sort purified from the thymus (CD4^+^CD8^+^, GFP^hi^) or spleen (GFP^med-hi^) of adult F1 mice generated from *Luc*^*shRNA*^. Vav-TA and *Kdm6b*^*shRNA*^. Vav-TA crosses and probed for *L32* and *Kdm6b* (d). Data shown are mean+sem and are representative of 1–6 mice, *p<0.05, ***p<0.001.

Having validated Dox-inducible *Kdm6b* knockdown in *Kdm6b*^*shRNA*^.CAG-rtTA bitransgenic MEFs, we then wanted to determine whether we could knockdown *Kdm6b* transcription within hematopoietic cells by generating *Kdm6b*^*shRNA*^.Vav-tTA transgenic mice, where the shRNA is expressed in the absence of Dox. To assess *Kdm6b* knockdown in developing T cells, we sorted purified GFP^hi^ CD4^+^CD8^+^ thymocytes, or GFP^med-hi^ (due to low cell number) splenocyte subsets from either *Luc*^*shRNA*^.Vav-tTA or *Kdm6b*^*shRNA*^.Vav-tTA mice (**Fig. 2d**). We observed a high level of *Kdm6b* knockdown within DP thymocytes with also substantial knockdown observed in naive CD4^+^ T cells (**Fig. 2d**). A modest level of *Kdm6b* knockdown (~30%) was observed in resting naive, or activated CD8+ T cells, while B cells and F4/80+ myeloid cells demonstrated low levels of knockdown, albeit from a low level of Kdm6b to start (**Fig. 2d**).

### Assessment of immature T cell commitment in Kdm6bshRNA.Vav-tTA transgenic mice

Expression of *Kdm6b* shRNA in untreated *Kdm6b*^*shRNA*^.Vav-tTA F1 offspring enabled the characterisation of *KDM6b* knockdown during T cell development in the thymus. High levels of *Kdm6b* shRNA expression (GFP^hi^) resulted in a lower frequency of CD4^−^CD8^−^ (DN), CD4^+^CD8^+^ (DP), CD4^+^ and CD8^+^ single positive thymocytes compared to those containing the *Luc* shRNA (**Fig. 3a, b**). This indicates that *Kdm6b* knockdown impacts thymocyte development. Overall cell numbers were not overtly impacted due to an increase in the frequency of GFP^med^ cells in *Kdm6b*^*shRNA*^ mice, compared to *Luc*^*shRNA*^ mice in these subsets (**Fig. 3a, b**).

**Figure 3.**
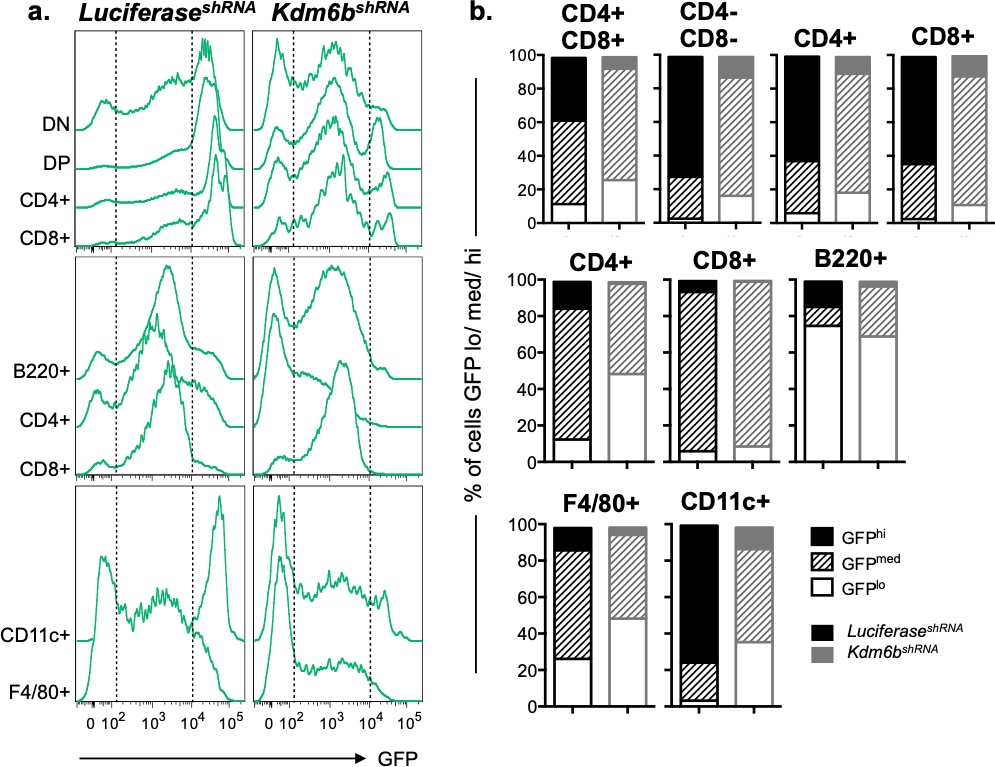
Expression levels of target hairpin in hematopoetic cells in *Luc*^*shRNA*^ and *Kdm6b*^*shRNA*^ mice. **(a)** GFP reporter expression in thymic CD4^−^CD8^−^ (DN), CD4^+^CD8^+^ (DP), or single CD4^+^ or CD8^+^ T cell subsets (upper panel) and splenic B220+ (B cells), naive CD4^+^ (CD4 T cells) and naive CD8^+^ (CD8 T cells) cells (mid panel), F4/80+ (macrophages) and CD11c+ (dendritic cells) cells (lower panel). **(b)** Quantification of proportions of cell subsets. Data shown are representative of 1-2 mice/strain and 3 independent experiments.

Analysis of mature, peripheral splenic cell subsets displayed a similar yet more extreme trend, whereby there are almost no GFP^hi^ expressing CD4^+^ or CD8^+^ T cells or B220+ B cells from mice bearing the *Kdm6b* hairpin, compared to the *Luc*^*shRNA*^ control (**Fig. 3a, b**, middle panels). For at least the T cell subsets, the proportion of GFP^hi^CD4+ T cells was lower in the *Kdm6b*^*shRNA*^ vs *Luc*^*shRNA*^ mice. There were few GFP^hi^CD8+ T cells with most cells expressing an intermediate level of GFP (GFP^med^) in both lines of mice. This suggests that there is a threshold effect in the T cell lineage whereby a low to intermediate level of shRNA transcription is not sufficient for *Kdm6b* knockdown. A similar trend is displayed in innate F4/80+ (macrophages; Mϕ) and CD11c+ (dendritic cells; DC) cell subsets between *Kdm6b* and *Luciferase* hairpin controls (**Fig. 3a, b**, lower panels) with selective outgrowth of GFP^lo^ Mϕs bearing the *Kdm6b* hairpin, compared to the *Luciferase* control. Taken together, these data suggest that constitutive expression of the *Kdm6b* shRNA impacts cellular development of hematopoetic cells. This is particularly apparent in thymic subsets and CD4^+^ splenocytes.

### Kdm6b knockdown impairs T cell development in the thymus

To further explore the impact of shRNA on T cell development, we assessed the proportion of DN, DP, CD4^+^ and CD8^+^ T cells in GFP^lo^ (shRNA^lo^) or GFP^hi^ (shRNA^hi^) subsets from the thymus of *Luciferase* or *Kdm6b* mice. While there was no general impact of *Kdm6b*^*shRNA*^ expression on the proportion of DP or SP thymocyte subsets, there was a consistent decrease in the frequency of DN cells in mice expressing either the *Luciferase* or *Kdm6b* hairpin indicative of a non-specific impact on DN thymocyte differentiation (**Fig. 4a**). To further delineate the differentiation stage being impacted, we analysed DN subsets (DN1; CD44+CD25-, DN2; CD44+CD25+, DN3; CD44-CD25+, DN4; CD44-CD25-; Ref. (29)) to determine if there was a specific role for KDM6b in T cell selection. In this case, GFP^hi^ thymocytes from *Kdm6b*^*shRNA*^ mice have a higher frequency of DN2 phenotype cells compared to *Luc*^*shRNA*^ control mice (**Fig. 4b**). The frequency of DN1, DN3 and DN4 subsets were comparable between *Kdm6b*^*shRNA*^ and *Luc*^*shRNA*^ control mice (**Fig. 4b**). Therefore, it is possible that KDM6b plays a role in the transition from the DN1 to DN2 phase of T cell development in the thymus.

**Figure 4.**
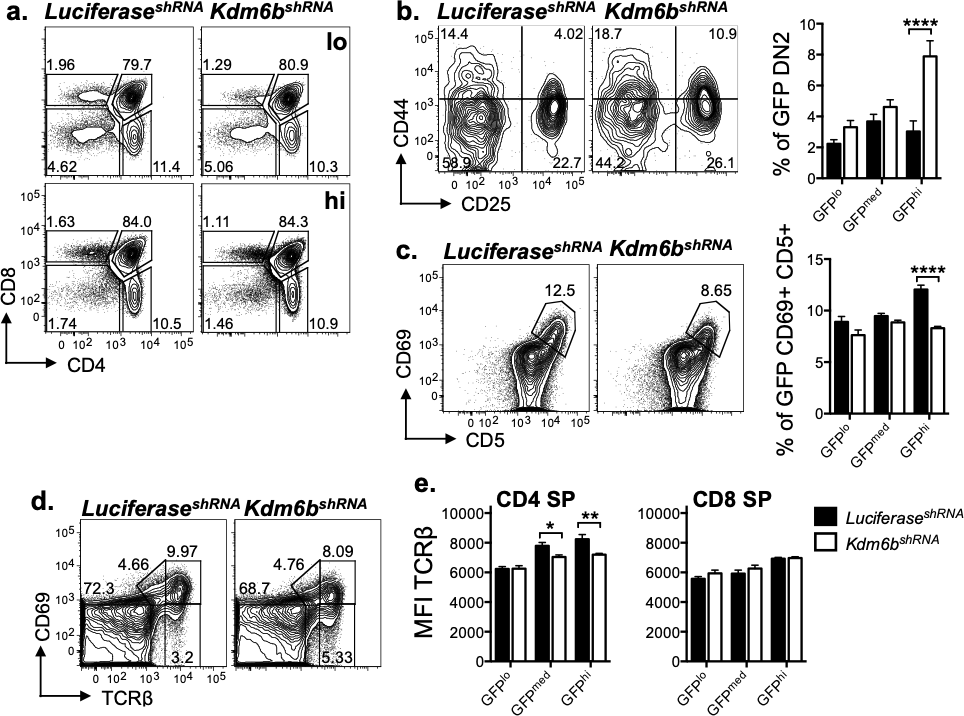
Determining the role of KDM6b during T cell development in the thymus. Flow cytometry profiles of CD4 versus CD8 subsets gated on GFP^lo^ (upper panel) or GFP^hi^ (lower panel) subsets from the thymus of *Luc*^*shRNA*^.Vav-tTA and *Kdm6b*^*shRNA*^.Vav-tTA mice (a). Flow cytometry profiles of DN1 (CD44+ CD25-), DN2 (CD44+ CD25+), DN3 (CD44-CD25+) and DN4 (CD44-CD25-) thymic subsets gated on GFP^hi^ cells from mice indicated in a. and quantified for increasing GFP expression (b). Flow cytometry profiles of CD69 versus CD5 subsets gated on GFP^hi^ cells from mice indicated in a. and quantified for increasing GFP expression (c). Flow cytometry profiles of CD69 and TCRβ thymic subsets gated on GFP^hi^ cells from mice indicated in a. (d). Mean fluorescent intensity (MFI) of TCRβ within TCRβ+ GFP^hi^ CD4^+^ (left) or TCRβ+ GFP^hi^ CD8^+^ (right) cells and quantified for increasing GFP expression (e). Data shown are mean+sem (n=4) are representative of four (a-b) or a single (c-e) experiment/s, *p<0.05, **p<0.01, ****p<0.0001.

To determine if KDM6b may also be impacting on T cell positive selection, we compared expression of CD69 and CD5 on GFP^hi^ thymocytes between *Kdm6b*^*shRNA*^ and *Luc*^*shRNA*^ control mice (**Fig. 4c**). There was a small but significant decrease in the frequency of CD69^+^CD5^+^ cells expressing the *Kdm6b* shRNA compared to the Luciferase control (**Fig. 4c**). These data point to a potential role of Kdm6b in either promoting positive selection, or alternatively, a lack of Kdm6b function may result in an increased sensitivity to negative selection. To further assess whether this effect was a defect in positive selection in *Kdm6b*^*shRNA*^ mice, we analysed the expression of TCRβ and CD69 on GFP^hi^ thymocytes. There was no significant difference in any of the CD69+TCRβ^int/hi^ subsets, suggesting that a difference in positive selection is likely to be subtle between *Kdm6b*^*shRNA*^ and *Luc*^*shRNA*^ control mice (**Fig. 4d**) (30). Finally, we analysed TCRβ MFI on CD4^+^ and CD8^+^ single positive cells between *Kdm6b*^*shRNA*^ and *Luc*^*shRNA*^ control mice (**Fig. 4e**). When graded for GFP expression, it is clear that with increased GFP, and therefore increased shRNA transcription, there was decreased TCRβ fluorescent intensity on CD4^+^, but not CD8^+^ single positive cells harbouring the *Kdm6b* shRNA (**Fig. 4e**). This suggests that Kdm6b knockdown may increase sensitivity to negative selection. Taken together, this further supports the notion that KDM6b may play a role in fine tuning sensitivity of thymocytes.

### KDM6b is involved in the maintenance of CD4^+^ T cells in the periphery

The impact of *Kdm6b* knockdown on SP CD4+ T cells in thymus lead us to ask whether there was also an effect on mature, peripheral lymphocytes. We first assessed the proportion of CD4^+^ (CD4 T cells), CD8^+^ (CD8 T cells), B220^+^ (B cells), NK1.1^+^ (NK cells), CD11c^+^ (dendritic cells) and F4/80^+^ (macrophages/monocytes) within naïve *Luc*^*shRNA*^.Vav-tTA and *Kdm6b*^*shRNA*^.Vav-tTA mice. There was a decrease in the frequency of GFP^hi^ CD4^+^ T cells in *Kdm6b*^*shRNA*^ transgenic mice (**Fig. 5a, b**). This appeared to be compensated for by an increase in the frequency of GFP^lo^ CD4^+^ T cells in *Kdm6b*^*shRNA*^ transgenic mice (**Fig. 5a, b**). Compared to CD4+ T cells, there was a much lower frequency of GFP^hi^ CD8^+^ T cells in both the *Kdm6b*^*shRNA*^ transgenic and *Luc*^*shRNA*^ control mice (**Figure 5a, b**). It was interesting that an increase in the frequency of GFP^med^ CD8^+^ T cells in *Kdm6b*^*shRNA*^ transgenic mice was observed compared to *Luc*^*shRNA*^ control mice (**Fig. 5b**). Hence, it is possible that KDM6b knockdown actually supports mature CD8+ T cell differentiation.

**Figure 5.**
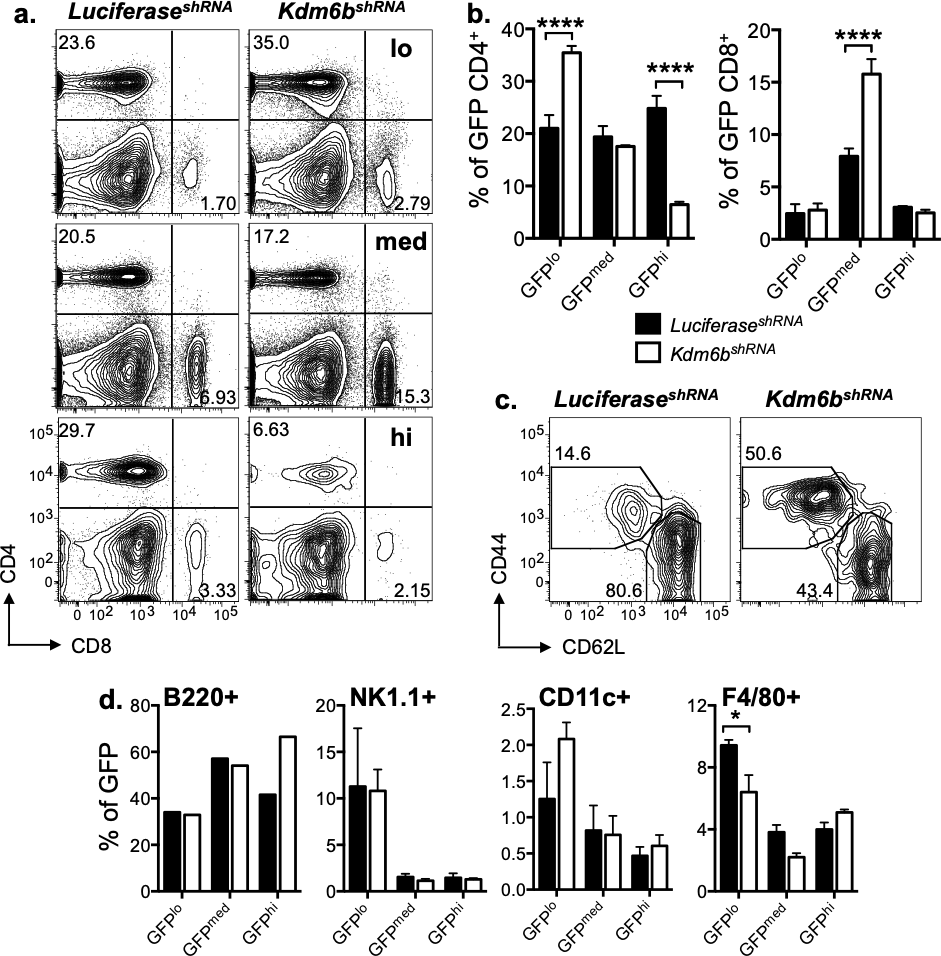
Determining the role of KDM6b in immune cell subsets in naïve mice. Flow cytometry profiles of CD4 versus CD8 subsets gated on GFP^lo^ (upper panel), GFP^med^ (middle panel) or GFP^hi^ (lower panel) subsets from the spleen of *Luc*^*shRNA*^.Vav-tTA and *Kdm6b*^*shRNA*^.Vav-tTA (a) and quantified for increasing GFP expression (b). Flow cytometry profiles displaying activation phenotype of CD4^+^ T cells gated on GFP^hi^ cells from mice indicated in a. (c). Frequency of splenic cells from indicated mice that are B220+ (B cells), NK1.1+ (NK cells), CD11c+ (dendritic cells) or F4/80+ (macrophages) quantified for increasing GFP expression (d). Data shown are mean+sem (n=4) and are representative of four (a-c), one/two (d) independent experiment/s, *p<0.05, ****p<0.0001.

Given that KDM6b knockdown within the GFP^hi^ expressing subsets resulted in a lower proportion of splenic CD4^+^ T cells, the phenotype of GFP^hi^ CD4^+^ T cells was analysed (**Fig. 5c**). Despite a lower proportion of total GFP^hi^ CD4+ T cells within *Kdm6b*^*shRNA*^ mice, there was an increase in the proportion of these cells exhibiting an activated phenotype (CD44^+^ CD62L^lo^) when compared to those expressing a high level of the *Luciferase* control hairpin (**Fig. 5c**). Taken together, these data indicate that KDM6b dysregulation in steady state appears to impact mature, naive CD4^+^ T cell development and persistence. KDM6b knockdown did not impact the proportion of splenic NK cells (NK1.1+), dendritic cells (CD11c+) or macrophages/monocytes (F4/80+) compared to the Luciferase control hairpin (**Fig. 5d**). Interestingly, there was an increased proportion of GFP^hi^ B220+ B cells in the spleens of *Kdm6b*^*shRNA*^ compared to *Luc*^*shRNA*^ transgenic mice (**Fig. 5d**).

### KDM6b knockdown results in dysregulation of an influenza-specific T cell response

While no impact on the proportion of naïve CD8^+^ T cells was observed, the initial hypothesis was that KDM6b would play a role in CD8^+^ T cell differentiation, via the resolution of key transcription factor loci that mediate the lineage specific function of these cells (16). Therefore, we wished to further test this hypothesis following infection of *Luc*^*shRNA*^.Vav-tTA and *Kdm6b*^*shRNA*^. Vav-tTA mice with influenza A virus (IAV).

Naïve *Luc*^*shRNA*^ and *Kdm6b*^*shRNA*^ mice were infected with 1×10^4^ plaque forming units (pfu) of A/HKx31 and weight loss measured over ten days (**Fig. 6a**). Both groups of mice lost a comparable amount of weight following infection and recovered fully, indicating no gross defects in viral clearance and pathology. As is seen in the naïve state, high expression of the *Kdm6b* shRNA in the spleens resulted in a dramatic decrease in the frequency of CD4^+^ T cells at the peak of the immune response (**Fig. 6b, c**), with no significant difference in the proportion CD8^+^ T cells between the *Luc*^*shRNA*^ and *Kdm6b*^*shRNA*^ transgenic mice (**Fig. 6b**). The preliminary trend seen for an increase in B220^+^ B cells in the presence of the Kdm6b hairpin in the naïve state remained consistent following infection (**Fig. 6b, c**). This is interesting given recent data showing a role for H3K27me3 in regulating germinal B cell responses (31). As seen in the naïve state, CD4^+^ T cells harbouring a high amount of the *Kdm6b* hairpin had an associated higher frequency of activated CD4^+^ T cells compared to those harbouring the *Luciferase* hairpin (**Fig. 6d, e**) suggesting that KDM6b knockdown somehow enabled CD4+ T cell differentiation.

**Figure 6.**
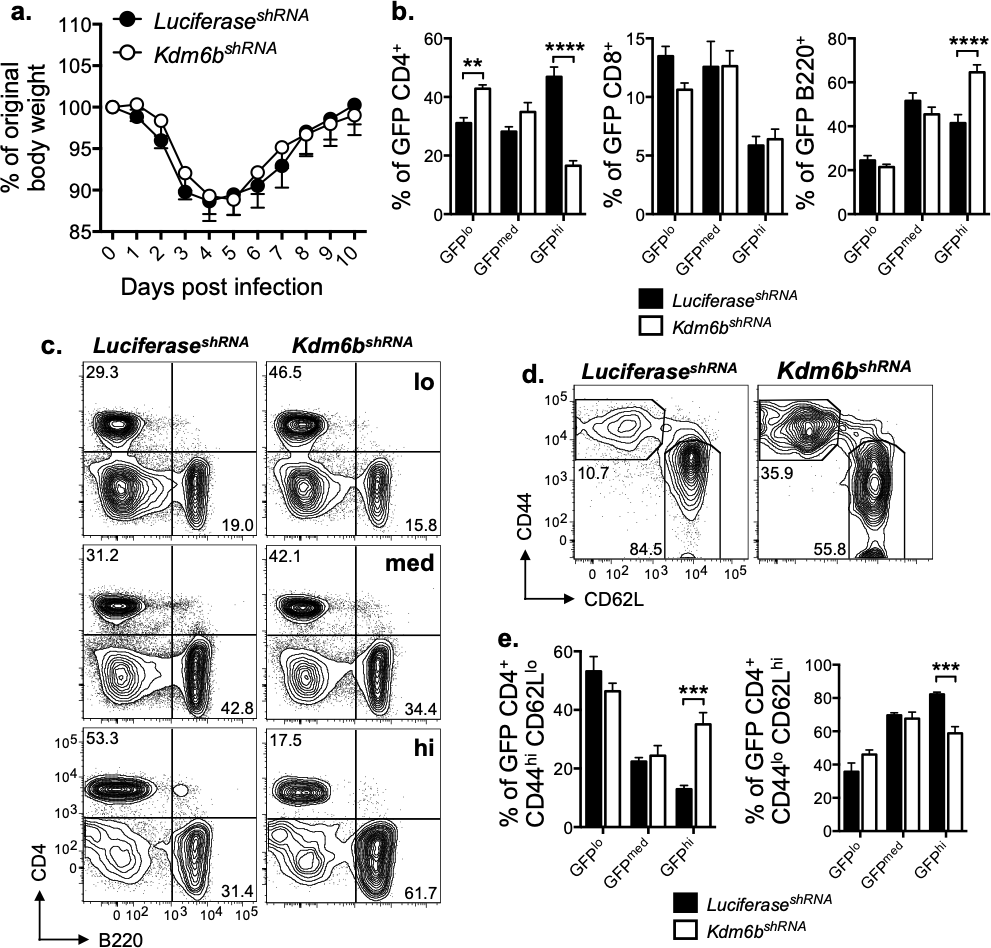
Determining the role of KDM6b in immune cell subsets in influenza A infected mice. Naïve *Luc*^*shRNA*^.Vav-tTA and *Kdm6b*^*shRNA*^.Vav-tTA mice were infected with X31 and weight loss measured over 10 d (a). Frequency of CD4^+^, CD8^+^ and B220^+^ cells was quantified for increasing GFP expression in (b) and representative flow cytometry profiles of CD4 versus B220 subsets gated on GFP^lo^ (upper panel), GFP^med^ (middle panel) or GFP^hi^ (lower panel) subsets from the spleen of *Luc*^*shRNA*^ and *Kdm6b*^*shRNA*^ mice are shown in (c). Flow cytometry profiles displaying activation phenotype of CD4^+^ T cells gated on GFP^hi^ cells from mice indicated in b. (d) and quantified for increasing GFP expression in (e). Data shown are mean+sem (n=5) and are representative of a single experiment, **p<0.01, ***p<0.001, ****p<0.0001.

To further investigate whether KDM6B knockdown impacted antigen-specific CD8^+^ and CD4^+^ T cell responses, the proportion of the CD8^+^ D^b^NP_366_ and D^b^PA_224_-specific and CD4^+^ I-A^b^/HA_211_-specific responses were determined after peptide stimulation and ICS for IFNγ within the GFP- or GFP+ (GFP^med-hi^) subsets (**Fig. 7**). However, there was a significant reduction in both the frequency and total number of D^b^NP_366_ and D^b^PA_224_ specific CD8^+^ T cells in GFP^+^ populations within the *Kdm6b*^*shRNA*^ mice, compared to the *Luc*^*shRNA*^ controls (**Fig. 7a-c**). Surprisingly, this trend was the opposite for I-A^b^/HA_211_-specific CD4^+^ T cells which displayed an increase in frequency compared to the *Luc*^*shRNA*^ controls but no difference in total cell number (**Fig. 7a-c**). There was no difference in the frequency of either IAV-specific CD8^+^ and CD4^+^ T cell responses, within the GFP^−^ populations from *Luc*^*shRNA*^ and *Kdm6b*^*shRNA*^ transgenic mice (**Fig. 7**) indicating the impact was due to *Kdm6b* shRNA expression. Hence, these data suggest that KDM6b may in fact play distinct roles in either IAV-specific CD8^+^ vs CD4^+^ T cell responses.

**Figure 7.**
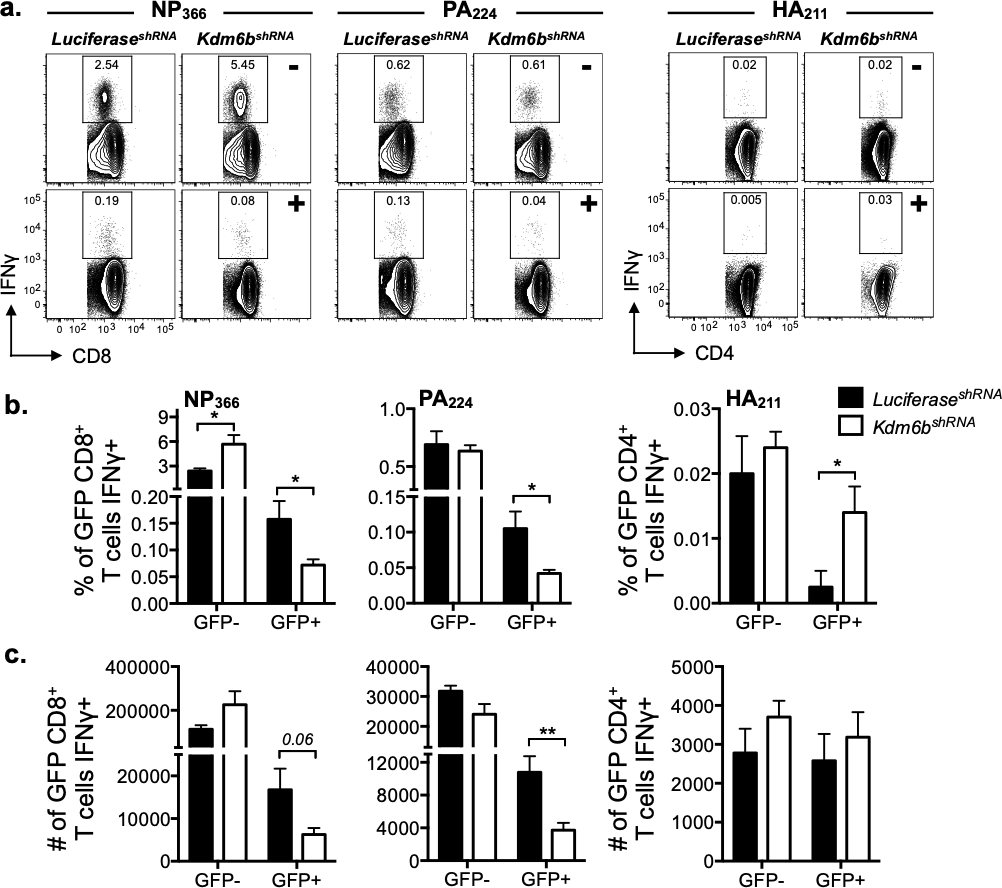
Determining the role of KDM6b in antigenspecific CD4^+^ and CD8^+^ T cells in IAV infected mice. Naïve *Luc*^*shRNA*^.Vav-tTA and *Kdm6b*^*shRNA*^.Vav-tTA mice were infected with X31 and 10 d post infection, the expression of IFNγ was determined by ICS ex vivo (a). Shown are representative flow cytometry profiles of CD8 or CD4 versus IFNγ gated on GFP-(upper panel) or GFP+ (lower panel) subsets from the spleen of *Luc*^*shRNA*^ and *Kdm6b*^*shRNA*^ mice and quantified for frequency (b) or total cell number (c). Data shown are mean+sem (n=5) and are representative of a single experiment, *p<0.05, **p<0.01.

### KDM6b may play a role in resolving bivalency of key transcription factor loci

Further, we have previously shown that in the naive state, the *Tbx21* locus in CD8^+^ T cells is marked with the permissive H3K4me3 and repressive H3K27me3 histone PTMs. Upon naive CD8+ T cell activation, the *Tbx21* locus is rapidly resolved to a permissive state following activation via loss of the KDM6b substrate H3K27me3 (16). To assess whether KDM6b expression played a role in regulating this resolution, we analysed the epigenetic signature of key transcription factors important for CD8^+^ T cell differentiation in the presence of the KDM6b hairpin at the peak of the immune response to IAV.

Naïve *Luc*^*shRNA*^ and *Kdm6b*^*shRNA*^ transgenic mice were infected with A/HKx31 and ten days post infection, cells were harvested from the spleens of *Luc*^*shRNA*^ and *Kdm6b*^*shRNA*^ mice and analysed for T-bet protein expression directly *ex vivo* by ICS on activated (CD44^hi^) CD4^+^ and CD8^+^ T cells (**Fig. 8**). This analysis was performed in order to determine if resolution of bivalency was likely to have occurred prior to performing epigenetic analysis. In this case, there was no significant decrease in the level of T-bet protein in activated CD4^+^ and CD8^+^ T cells following IAV infection of *Luc*^*shRNA*^ and *Kdm6b*^*shRNA*^ mice (**Fig. 8a**), despite an overall lower, albeit modest, level of *Kdm6b* RNA (**Fig. 8b**). Hence, perhaps unsurprisingly, there was little difference in the expression of transcription factor genes with bivalent loci in activated, polyclonal CD4^+^ or CD8^+^ T cells from either mouse strain (**Fig. 8c**).

**Figure 8.**
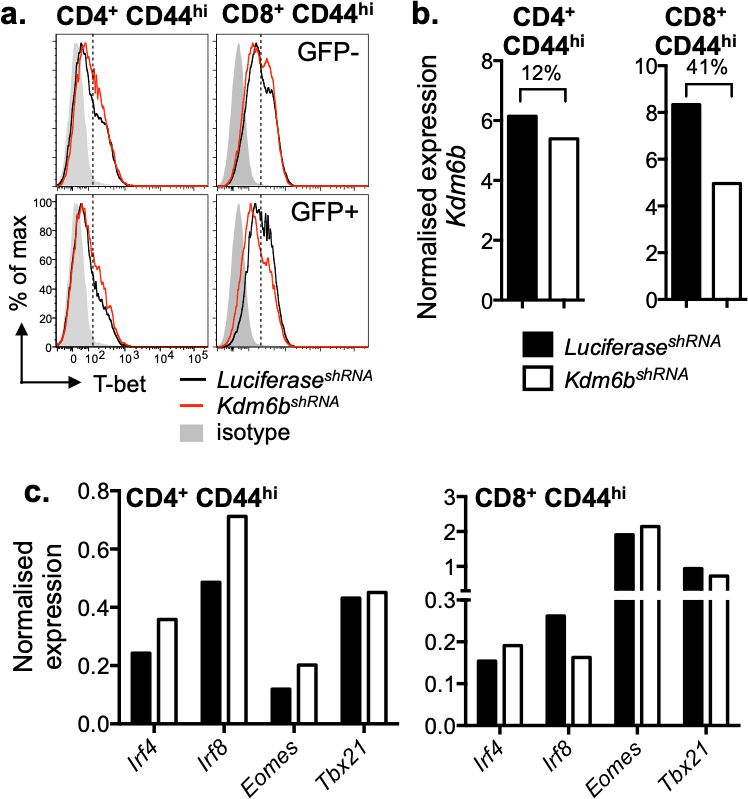
Determining the role of KDM6b in the regulating the expression of fundamental transcription factor loci following IAV infection. Naïve *Luc*^*shRNA*^.Vav-tTA and *Kdm6b*^*shRNA*^.Vav-tTA mice were infected with X31 and 10 d post infection, the expression of T-bet was determined by ICS *ex vivo*. Representative histograms show T-bet expression gated on indicated cells from GFP− (upper panel) or GFP+ (lower panel) subsets from the spleen of *Luc*^*shRNA*^ and *Kdm6b*^*shRNA*^ mice (a). GFP+ subsets were sort purified and probed for *L32* and *Kdm6b* (b) or *Irf4*, *Irf8*, *Eomes* and *Tbx21* (c). Data shown are from one experiment with n=5 mice/genotype.

Despite no overall difference in the levels of T-BET, we still examined whether *Kdm6b* knockdown could impact removal of H3K27me3 at specific gene loci. To test this, sort purified GFP^med-hi^ CD44^hi^ CD4^+^ and CD8^+^ T cells were isolated 10 days after IAV infection and chromatin immunoprecipitation (ChIP) carried out for the permissive modification H3K4me3 and the repressive modification H3K27me3 at bivalent gene loci (16). Interestingly, there was a trend for a greater level of H3K27me3 enrichment in the presence of the KDM6b hairpin at the *Tbx21* locus in activated GFPM^med-hi^ CD8^+^ T cells (**Fig. 9a**). Importantly, this same trend was not observed at the *Tbx21* loci in activated GFP^med-hi^ CD4^+^ T cells,nor at the loci of other transcription factors analysed in both CD8^+^ and CD4^+^(**Fig. 9a**). Interestingly, in activated GFP^med-hi^ CD8^+^ T cells there was an increase in the active modification H3K4me3 at *Ifr4* and *Irf8* loci, and suggesting a possible mechanism for the similar transcription levels between cells with the *Luciferase* and *Kdm6b* hairpin, despite the lack of difference in deposition of the repressive H3K27me3 (**Fig. 7a, b**). However, this trend was not consistent for the T-box transcription factors *Tbx21* and *Eomes* (**Fig. 7b**). Therefore, it remains unclear why the CD8^+^ T cells from mice harbouring the *Kdm6b* hairpin are able to express the T-bet protein and RNA despite the locus remaining unresolved in its chromatin state. It is possible that perhaps given that ChIP is a bulk assay, we are unable to discriminate those cells expressing T-BET that have lost H3K27me3. Further, it is important to note that the unresolved *Tbx21* loci in the presence of the *Kdm6b* hairpin does not appear to be a general mechanism as it is not seen in CD4^+^ T cells. Finally, the *actin* loci is used as a control and is expected to have constitutive deposition of H3K4me3 and lack H3K27me3. In the presence of the *Kdm6b* hairpin we see a decrease in the deposition of H3K4me3 at this locus possibly suggesting a general decrease in accessibility of chromatin in cells harbouring the *Kdm6b* hairpin.

**Figure 9.**
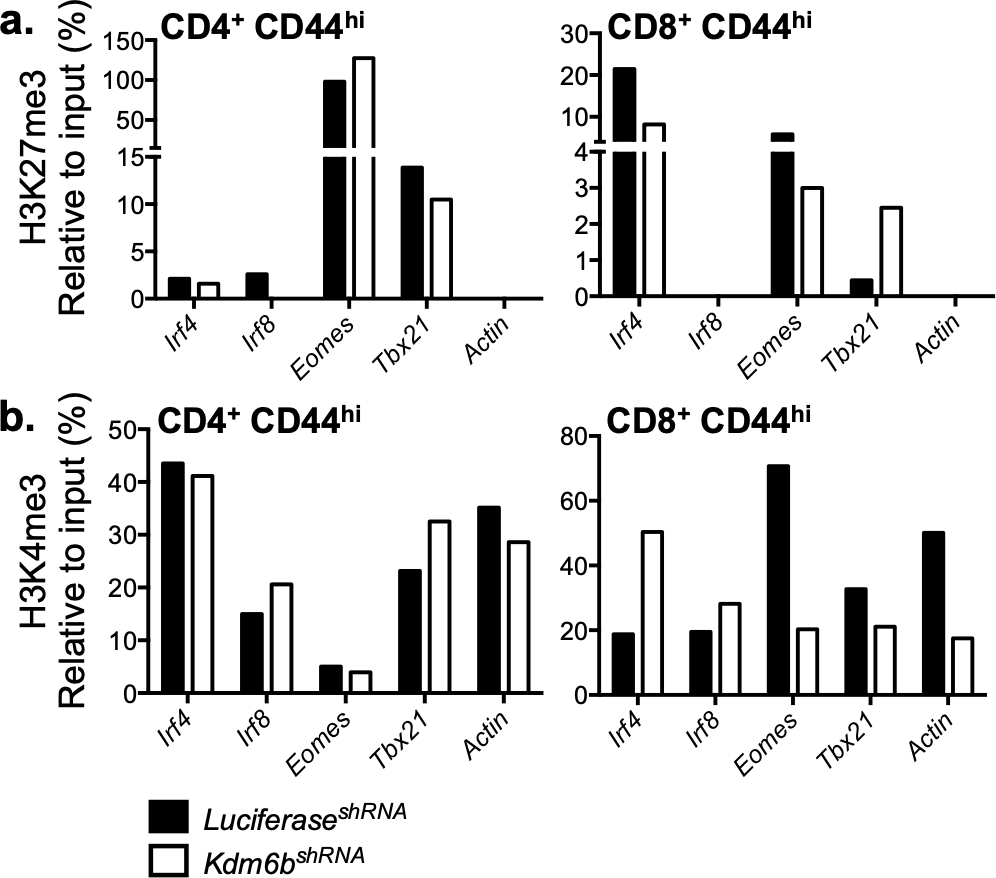
Determining the role of KDM6b in the resolution of bivalency in fundamental transcription factor loci following IAV infection. Naïve *Luc*^*shRNA*^.Vav-tTA and *Kdm6b*^*shRNA*^.Vav-tTA mice were infected with X31. Primers were designed to span the promoter region of indicated loci and 10 d post infection, GFP+ CD44^hi^ CD8^+^ and CD4^+^ T cells were sort purified, fixed at 0.6% and analysed by ChIP for deposition of H3K27me3 (a) or H3K4me3 (b). Analysis was conducted by qPCR, Ct value was converted to copy number, no antibody control subtracted and normalised to input DNA. Data shown are from one experiment with n=5 mice/genotype.

## Discussion

The use of RNAi mediated gene silencing induced by shRNA expression within a microRNA backbone is a potent way to decrease target gene expression (20). The utility of this system was demonstrated in an early study by Dickins *et al*. demonstrating that induction of shRNA knockdown targeting the tumour suppressor, p53, resulted in the development of tumours, with restoration of p53 expression resulting in decreased tumour burden (21). This use of tetracycline responsive regulatory element provides a powerful tool to be able to either turn on or turn off constitutive shRNA transcription. This study describes the development of two mouse lines designed to enable inducible and reversible hairpin mediated knockdown of the epigenetic modifier KDM6b.

We first generated mice where *Kdm6b* knockdown was induced upon addition of doxycycline to drinking water or food. By crossing *Kdm6b*^*shRNA*^ or *Luc*^*shRNA*^ transgenic mice with CAG-rtTA3 (Tet-ON) mice, robust shRNA expression upon doxycycline induction was evident in adaptive immune cell subsets (25). Using this system would enable shRNA expression to be induced at any stage during an immune response *in vivo*. Interestingly, this system appears to function well in developing thymocytes, yet some mature T cell subsets have been reported to be refractory to to shRNA transcription. Nevertheless, this system provides increased flexibility and precision in understanding when this epigenetic modifier plays a role in the programming of multiple immune cell types.

An advantage of the shRNA mediated knockdown system, compared to permanent gene mutation or removal is that the target gene is depleted instead of being completely deleted (22). In some cases this more accurately reflects human disease (24, 32). For example, it has been demonstrated that *Kdm6b* deletion can lead to embryonic lethality (33) or deformity (34, 35). Hence, the ability to control KDM6b knockdown via shRNA induction enables this issue to be overcome.

To that end, we established a line of mice using the “Tet-OFF” system whereby there is constitutive *Kdm6b* shRNA dependent knockdown that can be reversed by addition of doxycycline (*Kdm6b^shRNA^. Vav-tTA* mice). We were able to demonstrate *Kdm6b* knockdown in a variety of immune cell types including immature thymocytes and mature T cells. Our preliminary analysis of thymocyte development indicated that KDM6b may play a role in T cell development in the thymus, specifically during transition from the DN1-DN2 stage. Recently, Manna and colleagues have shown that both KDM6b and the related enzyme UTX are essential for terminal thymocyte differentiation, specifically due to their ability to remove the repressive H3K27me3 mark from genes associated with egress such as *S1pr1* (36). Dynamic changes in H3K27me3 deposition is a characteristic of thymocyte differentiation (37). This system therefore may be useful to further delineate the precise role of KDM6b in this process.

We observed, even with a relatively modest knockdown of *Kdm6b*, fewer mature CD4^+^ T cells in the periphery, both in the naïve state and following IAV infection. While the defect in the DN2 transition maybe an explanation, there was no impact on either DP or SP populations, and no impact on the mature CD8+ T cell numbers in the periphery making this unlikely. It was interesting to note that mature, naive CD4^+^ T cells in *Kdm6b^shRNA^* mice had a more activated phenotype, reminiscent of T cells that have undergone some level of activation or homeostatic proliferation, termed virtual memory T cells (T_VM_) (38). Hence, KDM6b expression maybe required for maintenance of a naive state within CD4^+^ T cells, perhaps by helping limit responsiveness to homeostatic cytokines such as interleukin-15, known to drive T_VM_ formation (39).

Interestingly, the decrease in the frequency of CD4^+^ T cells correlated with an increase in the frequency of B cells. An explanation as to why we see this phenotype was not directly tested but one hypothesis is the decrease in the frequency of CD4^+^ T cells likely resulted in an increase in IL-7 concentration in the spleen, which is seen in lymphopenic conditions enabling increased survival of other cell types (40). While the total CD4^+^ T cell pool is decreased in *Kdm6b*^*shRNA*^ mice, the relative contribution of particular TH subsets such as TH1 vs TH2 was not assessed. A recent publication using a *Kdm6b* conditional knock out mouse (*Kdm6b*^*fl/fl*^.*CD4*^*cre*^) showed that depletion of *Kdm6b* selectively promoted Th2 and Th17 differentiation while inhibiting Th1 differentiation (12). It remains possible that the lack of KDM6b may result in a skewing of CD4+ T cells into specific subsets such as T_H_2 or Follicular CD4+ T cells. In both cases, either the production of IL-4 by T_H_2 cells which can promote B cell survival and proliferation, and/or increased B cell proliferation due to greater germinal centre formation supported by T_FH_ cell may be an explanation (41). Importantly, an increase in the Tfh cell subset would also increase IL-21 production which has been linked to B cell proliferation and antibody secretion (42). Furthermore, a lack of *Kdm6b* prevented conversion of T_H_2 or T_H_17 CD4^+^ T cells into Treg cells. Hence, if Treg cells were selectively affected by a decrease in KDM6b expression, this could cause a further increase in CD4 subssets that could exacerbate expansion or maintenance of B cells.

While the proportion and number of mature, naive CD8^+^ T cells was not impacted in *Kdm6b^shRNA^* transgenic mice, upon IAV infection, fewer splenic D^b^NP_366_ and D^b^PA_224_ specific CD8^+^ T cells were observed at the peak of the IAV immune response. We have previously demonstrated that loss of H3K27me3 is a major consequence of virus-specific CD8^+^ T cell activation (15–17). Importantly, we identified a subset of gene promoters that exhibited bivalency for both H3K4me3 and H3K27me3. These loci marked transcription factors, such as *Tbx21*, known to play important roles in optimal virus-specific CD8+ T cell differentiation (43–45). Given these results, we speculated that KDM6b induction is a key step for optimal virus-specific CD8+ T cell differentiation. To that end, we observed that the bivalent *Tbx21* locus failed to resolve to a transcriptionally permissive state (H3K4me3+) in *Kdm6b*^*shRNA*^ activated CD8+ T cells. Interestingly, we saw little difference in the expression T-BET, and other key lineage specific transcription factors within IAV-specific T cells. Moreover, other bivalent loci such as *Irf4, Irf8*, and *Eomes* were able to be resolved to a permissive state. Furthermore, the *Tbx21* promoter was resolved in activated CD4^+^ T cells, indicating that KDM6b mediated resolution of bivalency at the *Tbx21* promoter was a CD8^+^ T cell specific mechanism. So while there is some hint that H3K27me3 removal from the *Tbx21* promoter is KDM6b dependent, and this correlates with a reduced virus-specific CD8+ T cell response, it remains to be determined whether this is a direct or indirect effect. Analysis of genome wide patterns of H3K27me3 deposition within both *Kdm6b*^*shRNA*^ and *Luc*^*shRNA*^ virus-specific T cells may provide more mechanistic information.

Another possibility for the subtle observations made is that there is a KDM6b homologue, KDM6a (UTX), that is also a H3K27 demethylase. Interestingly, it has been shown to be essential for embryonic development via a mechanism that is independent of its H3K27 demethylase activity (33). This demethylase independent function appears to be due to its the ability of UTX to bind the genome and recruit signalling factors (46, 47). Hence, the subtle differences observed may be due to overlapping or secondary roles of UTX in regulating T cell specific gene transcription.

An interesting note is that studies by Weinmann and colleagues have shown that KDM6b and UTX can interact with T-bet directly enabling TF regulation of TFs that is independent of H3K27 demethylase activity (48). It was demonstrated that physical interactions can occur between T-bet, Set7/9 H3K4 methyltransferases, and also the chromatin remodelling Brg1-containing SWI/SNF complex (13, 48, 49) were dependent on KDM6b, UTX, but did not require H3K27 demethylase activity. This provides a basis of how KDM6b can be recruited and co-locate to H3K4me3-marked loci and contribute to gene transcription and also how KDM6b is able to be recruited to target genes (50). Given the important role of T-bet in ensuring optimal virus-specific T cell responses, perhaps KDM6b knockdown is limiting the ability of T-bet to bind to target loci due to diminished KDM6b interactions. This would provide an explanation of why we see no significant difference in T-bet expression levels, yet observed diminished IAV-specific CD8+ T cell responses. In line with this possibility, we also observed decreased IFN-γ production, a known T-BET target gene, within IAV-specific CD8+ T cells. Analysis of T-bet binding to target gene loci within the *Kdm6b*^*shRNA*^ T cells via chromatin immunoprecipitation assays will provide further information about this potential mechanism.

## Acknowledgements

This work was supported by grants from the National Health and Medical Research Council of Australia (Program Grant #5671222 awarded to awarded to S.J.T; Project grant #APP1003131 (awarded to S.J.T). S.J.T is supported by an NHMRC Principal Research Fellowship. J.P. is supported by an Australian Postgraduate award. MEB was supported by a Bellberry Viertel Senior Medical Research Fellowship. Additional support was provided by the Victorian State Government Operational Infrastructure Support, Australian National Health and Medical Research Council IRIISS grant (9000433).

